# Genetic and structural interpretation of NLRP1 FIIND domain variants in glioma in an Indian cohort: a pilot study

**DOI:** 10.64898/2026.09.04.749492

**Authors:** Shalini Chhipa, Omkar A. Sonawane, M S Revanth, Deepak Kumar, Jyoti Sharma, Srijita Acharya, Deepak Jha, Sucharita Dey, Pankaj Yadav, Sushmita Jha

## Abstract

Gliomas, particularly glioblastoma (GBM), are aggressive primary brain tumours associated with dysregulated NLR signalling, a pathway central to innate immunity and inflammation. *NLRP1* triggers proinflammatory cytokine release and pyroptotic cell death via autoproteolytic cleavage. The FIIND missense variant rs11651270 (M1184V) may modulate this cleavage process. While *NLRP1* polymorphisms are associated with various diseases and cancers, their specific impact on glioma remains to be investigated. In our study, five FIIND-domain single-nucleotide polymorphisms (SNPs) of *NLRP1*-rs371579423, rs58604457, rs57636751, rs11651270, and rs2301583 were investigated by Sanger sequencing in a clinical cohort of glioma patients and compared against population-matched controls from the GenomeIndia dataset (Rajasthan cohort) using genetic association models. Molecular dynamics simulations were performed to evaluate the structural effects of the missense variant rs11651270 (M1184V) in *NLRP1* during pre- and post-cleavage states. The linked variants rs58604457 (G>A) and rs57636751 (C>T) exhibited complete linkage disequilibrium (r^2^=1.00) and were significantly associated with lower odds of glioma (Odds Ratio~0.5). The missense variant rs11651270 (T>C) showed no association with glioma risk. Notably, the SNP rs2301583 had a higher allelic frequency in glioma cases despite being absent in the GenomeIndia population-based control catalogue. Molecular dynamics simulations revealed that the M1184V substitution stabilizes local FIIND architecture by preserving β-strand organization through persistent interactions with neighbouring residues. This study provides the first combined genetic and structural analysis of *NLRP1* FIIND-domain variants in an Indian glioma cohort. These findings illustrate the potential value of integrating population-based genetic association with structural modelling to generate hypotheses and uncover potential functional mechanisms of inflammasome-gene variants in neuro-oncology.

**Highlights:**

- Two NLRP1 FIIND-domain SNPs are in complete linkage disequilibrium and associated with a lower odds ratio.
- The SNP rs2301583 exhibited an elevated allelic frequency in glioma cases, whereas it was not present in the GenomeIndia population-based control database.
- M1184V–E1195 hydrogen bonding locally stabilizes FIIND after cleavage.

## Introduction

Gliomas are the most prevalent primary central nervous system (CNS) tumours characterized by marked clinical heterogeneity, aggressive local invasion, and poor overall survival [1,2]. Glioblastoma (GBM), the most aggressive, malignant form, carries a median overall survival of approximately 14 months [3]. The fifth edition of the WHO classification of CNS tumours emphasizes an integrated diagnostic framework combining histopathological features with molecular markers such as IDH1/2 mutation status and 1p/19q codeletion to refine diagnostic grading, predict clinical behaviour, and guide therapy [4,5]. Tumour-associated genetic alterations modulate the expression of inflammatory mediators, recruitment of inflammatory cells, and shape the tumour microenvironment [6]. Chronic neuroinflammation is increasingly recognized as a key driver of glioma initiation, progression, and therapeutic resistance [7,8]. Epidemiological and preclinical evidence suggest a link between chronic neuroinflammatory states and a possible increase in glioma risk [9].

Inflammatory signalling within the glioma microenvironment is initiated by pattern-recognition receptors (PRRs) which sense pathogen- and damage-associated molecular patterns (PAMPs and DAMPs) to regulate downstream cytokine release and immune cell recruitment [10–12]. Among these, the nucleotide-binding domain leucine-rich repeat (NLR) family plays a central role in innate immunity. *NLRP1* (NLR family pyrin domain containing 1), responds to diverse PAMPs [13] such as anthrax lethal toxin, *Toxoplasma gondii,* and and metabolic stress (depletion of ATP) by undergoing autocatalytic cleavage to assemble into a multi-protein complex referred to as the *NLRP1* inflammasome [14], that drives the cleavage of enzymes and regulates biological processes such as inflammation, cellular homeostasis, metabolism, cell death [15–17]. Structurally, human *NLRP1* contains an N-terminal pyrin domain, nucleotide-binding (NACHT) domain, leucine-rich repeats (LRRs), a function-to-FIIND domain (FIIND), and a C-terminal caspase recruitment (CARD) domain [18]. Autoproteolytic cleavage within the FIIND domain cleaves *NLRP1* at the N-terminus; the N-terminal fragment is then degraded, releasing the C-terminal region, which initiates the downstream pathway [19]. However, enhanced FIIND autoproteolysis does not necessarily translate into enhanced inflammasome activation, because *NLRP1* activity is also regulated by the stability and turnover of mature *NLRP1*.

Genetic variation within the FIIND domain can alter cleavage kinetics, domain conformation, molecular dynamics, and downstream signalling capacity [20]. A missense variant caused by the single-nucleotide polymorphism (SNP) rs11651270 lies within the FIIND domain and leads to a mutation from methionine 1184 to valine (M1184V) and has been associated with variable disease phenotypes, including asthma susceptibility [21]. Structural characterization of the isolated, uncleaved FIIND domain by Moecking et al. demonstrated that the M1184V substitution reduces conformational flexibility around the autoproteolytic catalytic triad relative to methionine, thereby stabilizing the domain and promoting an autoinhibited *NLRP1*-DPP9 complex [22]. However, these analyses were restricted to the pre-cleavage state of the FIIND, and the structural consequences of M1184V on the post-cleavage FIIND– CARD domain interrelationship remain uncharacterized. Furthermore, the relevance of this variant in neuro-oncology has not been evaluated.

Although *NLRP1* polymorphisms, in coding and non-coding regions, have been associated with several diseases, including cancer, autoinflammatory, autoimmune, and neurological disorders [23], the contribution of *NLRP1* germline variation to glioma susceptibility remains largely unexplored. Initial inspection of publicly available data from The Cancer Genome Atlas (TCGA) [24] indicated a low overall burden of somatic *NLRP1* alterations in gliomas. However, low somatic mutation frequencies do not preclude a role for germline variants in modulating disease risk. Given the distinct genetic architecture across diverse human populations, we hypothesized that population-specific *NLRP1* variants contribute to glioma susceptibility in South Asian populations.

To test this hypothesis, we leveraged whole-genome sequencing data from the GenomeIndia initiative, a population-scale catalogue capturing genetic diversity across 10,000 individuals [25]. Using a geographically matched control cohort from the Rajasthan subset of GenomeIndia, we identified five FIIND-domain single-nucleotide polymorphisms (SNPs) in an Indian glioma cohort using Sanger sequencing and performed formal genetic association testing for four variants (rs2301583 was excluded from comparative association analysis as it was absent from the reference catalogue). To complement these genetic findings, we performed molecular dynamics simulations using AlphaFold2-generated models to evaluate the structural effects of the rs11651270 (M1184V) variant on protein stability and post-cleavage FIIND–CARD domain interactions. To our knowledge, this study provides the first genetic evaluation of *NLRP1* variants in an Indian glioma cohort and the first atomistic model of the post-cleavage FIIND–CARD interface for the M1184V substitution.

## Materials and Methods

### 1. Sample collection and DNA extraction

All samples in the study were from individuals diagnosed with glioma (grade 1 to 4), with ethical approval obtained. A total of 70 glioma tumour samples were used, including a male-to-female ratio of 2.5:1. Controls were extracted from the GenomeIndia panel’s Rajasthan subset, rather than age/sex-matched controls. Tumour samples were collected from AIIMS, Jodhpur, India, and preserved in RNA LivTM solution at 4°C, then stored at −20°C. Tissues were washed with 1X PBS prior to genomic DNA extraction using the Qiagen DNeasy Blood and Tissue kit, and DNA concentration and purity were measured with a NanoDrop™ 2000c Spectrophotometer.

### 2. PCR and Cleanup

A single 1.042 kb DNA fragment encompassing the *NLRP1* gene region was amplified using the TaKaRa Ex Taq® Hot Start Version kit. The PCR was set up with 10 μL of reaction mix using the final 1X Ex Taq Buffer, 0.2 mM dNTPs, 400 nM of each forward primer (5’– CGCACTTGGTCTTTGATGT–3’) and reverse primer (5’– AGAGGGTACCTGAGCCTAAG–3’), 0.25 U of TaKaRa Ex Taq HS enzyme, and template DNA. The unused/excess dNTPs and primers were cleaned up using ExoSAP-IT™ reagent (Applied Biosystems).

### 3. Sanger sequencing

Cycle sequencing was performed using the BigDye™ Terminator v3.1 Cycle Sequencing Kit (Applied Biosystems) as per the manufacturer’s instructions using the forward PCR amplification primer, followed by sodium acetate-EDTA cleanup. The final product was dissolved in Hi-Di™ Formamide (Applied Biosystems), followed by heat shock (95°C) for 5 min and snap-chilled (on ice) for 5 min before loading in an automated sequencer (SeqStudio Flex, Applied Biosystems) with a 36 cm length capillary and POP-7™ polymer (Applied Biosystems).

### 4. Construction of a population-specific allele frequency reference

A population-specific allele frequency reference was constructed using genomic data from the subset of the GenomeIndia data (Rajasthan population). The GenomeIndia subset was extracted from a larger genotype dataset and processed using PLINK v2.0[26]. Quality control filtering was applied prior to allele frequency estimation. Variants with missing genotype rates greater than 0% (--geno 0) were excluded to retain only fully genotyped markers. Samples with excessive missing genotypes (--mind 0.05) were removed to avoid biases arising from individuals with excessive missing genotypes. The variants with minor allele frequency <1% were removed to reduce the influence of rare genotyping artifacts. Variants deviating from Hardy–Weinberg equilibrium (HWE) (p < 1 × 10□□) were also filtered to remove markers potentially affected by genotyping artifacts or technical errors. After the HWE filter, heterozygosity rates were calculated for each individual. Samples showing extreme heterozygosity values (|F| > 0.1) were removed as potential outliers. After applying these filtering steps, allele frequencies were computed on the resulting processed dataset.

The resulting allele frequency file contains, for each variant, the chromosome and position, reference and alternate alleles, the alternate allele frequency, and the total number of observed chromosomes. This dataset provides a population-specific estimate of allele frequencies and serves as a population-specific germline reference comparable in concept to large-scale resources such as gnomAD, but calibrated for the local Rajasthan population.

### 5. Population frequency-based variant filtering

Variants identified in the tumour sequencing dataset were compared against allele frequencies observed in the Rajasthan dataset to distinguish common germline polymorphisms from potential tumour-associated variants. Population allele frequencies provide an important baseline for interpretation, particularly in targeted sequencing studies where the number of samples is limited.

Variants were interpreted according to their distribution across tumour samples and the population reference dataset. Variants observed at relatively high frequency in both the tumour dataset and the Rajasthan population were considered likely germline polymorphisms. Variants common in the Rajasthan population but detected at low frequency in tumour samples were interpreted as background population variants. In contrast, variants rare or absent in the Rajasthan population dataset but present at a higher frequency in tumour samples were considered potential tumour-associated variants. Variants rare in the population dataset but detected at moderate frequency in tumour samples were flagged as candidate variants requiring further validation.

### 6. Extraction of *NLRP1* variants from the population dataset

To enable direct comparison with the Sanger sequencing data, variants located within the *NLRP1* genomic locus were extracted from the Rajasthan population dataset. A slightly broader genomic interval surrounding the *NLRP1* gene was selected rather than restricting extraction strictly to the amplicon coordinates. This approach ensured that variants located near the boundaries of the sequenced region were not inadvertently excluded. Two reference VCF datasets were generated from this extraction, the first VCF dataset containing genotype information for 756 individuals and the second VCF dataset comprising a randomly selected subset of 70 individuals, generated using reproducible random sampling, provided that both datasets were adopted from the Rajasthan cohort. The smaller subset was created to facilitate exploratory analyses and computational testing with respect to tumour-representative population allele frequencies. By integrating tumour variant calls with population-specific allele frequency estimates, this approach provided an additional layer of filtering for distinguishing common germline variants from variants that may warrant further investigation in the context of glioma.

### 7. Genotype Quality Control and HWE

HWE was assessed in the control population (N=756). A chi-square test with one degree of freedom was used to test its statistical validity. Once the HWE of each rsID was confirmed, they were then subjected to further analysis (Supplementary Table S1)

### 8. Linkage Disequilibrium (LD) and Significance Thresholds

LD was calculated for all pairwise combinations of the four SNPs included in the case-control association using both r^2^ and D′ coefficients in both the case and control cohorts. Analysis revealed that rs58604457 and rs57636751 were in complete LD (r^2^ = 1.00, D′ = 1.00) (Supplementary Table S2), indicating that these two variants represented a single underlying association signal. The remaining variants, rs371579423 and rs11651270, showed distinct LD patterns. Thus, the four analysed SNPs represented three independent association signals: rs371579423, rs11651270, and the rs58604457/rs57636751 LD pair. The threshold for statistical significance was established at p < 0.05. All statistical analyses were performed using R (version 4.1) with base statistical packages. All the graphs were prepared using GraphPad Prism 10.0.0.

### 9. Statistical Analysis and Genetic Models

Genotype and allele frequencies for glioma cases and controls from the Rajasthan population were calculated, and Fisher’s exact tests were used to assess statistical significance. To evaluate the risk associated with specific variants, odds ratios (ORs) with corresponding 95% confidence intervals (CIs) were calculated.

Association analyses were performed using allelic and genotypic models, followed by three inheritance models: dominant, recessive, and codominant. For each analytical framework, all four SNPs included in the case-control analysis (rs371579423, rs11651270, rs58604457, and rs57636751) were evaluated individually. The dominant, recessive, and codominant inheritance models were analysed separately and were not included as an additional multiplicity factor in the Bonferroni correction. Bonferroni correction was applied across the three independent association signals identified by LD analysis within each analytical framework. The threshold for statistical significance was established at *p* < 0.05 after Bonferroni correction. Clinical associations for all SNPs were further explored by correlating genotypes with tumour grade, patient gender, and age using clinical data merged from glioma records.

### 10. Structure Modelling and Molecular Dynamics Simulations

The full structure of *NLRP1*, including FIIND and CARD domains, was not available in the PDB. We modelled the structure of the C-terminal fragment (residues 991-1473) using AlphaFold2[27]. The UniProt [28] sequence of the 483-residue fragment was used to obtain an initial model of the FIIND-CARD architecture encompassing the domains and the mutant (M1184V) and autoproteolytic cleavage (His 1186) sites. The highest-ranked model (ranking score = 0.71, pTM = 0.6) showed no steric clashes and moderate confidence in domain architecture and was used as a starting structure for Molecular Dynamics (MD) simulations. MD simulations were performed using GROMACS 2024[29] and the Amber99sb-ildn force field to investigate the effects of the M1184V mutant on global and FIIND-CARD inter-domain dynamics. We first simulated the wild-type and mutant full-length protein fragments to assess the pre-cleavage dynamics as affected by the mutant. Next, we removed the peptide bond at the cleavage site (His1186) to mimic the post-cleavage state and again simulated the cleaved structure for both wild-type and mutant.

Structural stability and conformational dynamics were characterized using multiple metrics, including backbone root-mean-square deviation (RMSD), radius of gyration (Rg), per-residue root-mean-square fluctuation (RMSF), solvent-accessible surface area (SASA) of the CARD domain, centre-of-mass distance between the FIIND and CARD domains, and hydrogen-bond analysis of both the cleavage pocket and the FIIND–CARD interface.

## Results

As previously mentioned, the analysis was carried out by comparing the cases with the population-scale catalogue and identified five SNPs. Interestingly, the fifth SNP rs2301583 was not found in the population-scale catalogue but was detected in the cases with an MAF of 6.0%, which is 4.6 times higher than its reported global MAF of 1.3% (Supplementary Table S3). Although the variant is currently annotated as benign in ClinVar, its relatively higher frequency indicates that it is worth exploring in the context of the Indian population.

### 1. Genotype Frequency Distribution of FIIND Domain Variants across the Study Cohort

Allele frequencies for all identified SNPs of *NLRP1* were compared between 70 glioma cases and the GenomeIndia subpopulation (N=756). For rs11651270, the heterozygous genotype frequencies were nearly identical between cases and controls across all subset comparisons, with odds ratios spanning between 0.716 and 2.13, suggesting no evidence of association. Interestingly, the minor alleles of rs58604457 and rs57636751 occurred at lower frequencies in cases than in controls, supporting the allelic and genotyping associations. Similarly, the rs371579423 allele showed no significant difference in frequency between cases and controls.

**Fig. 1.**
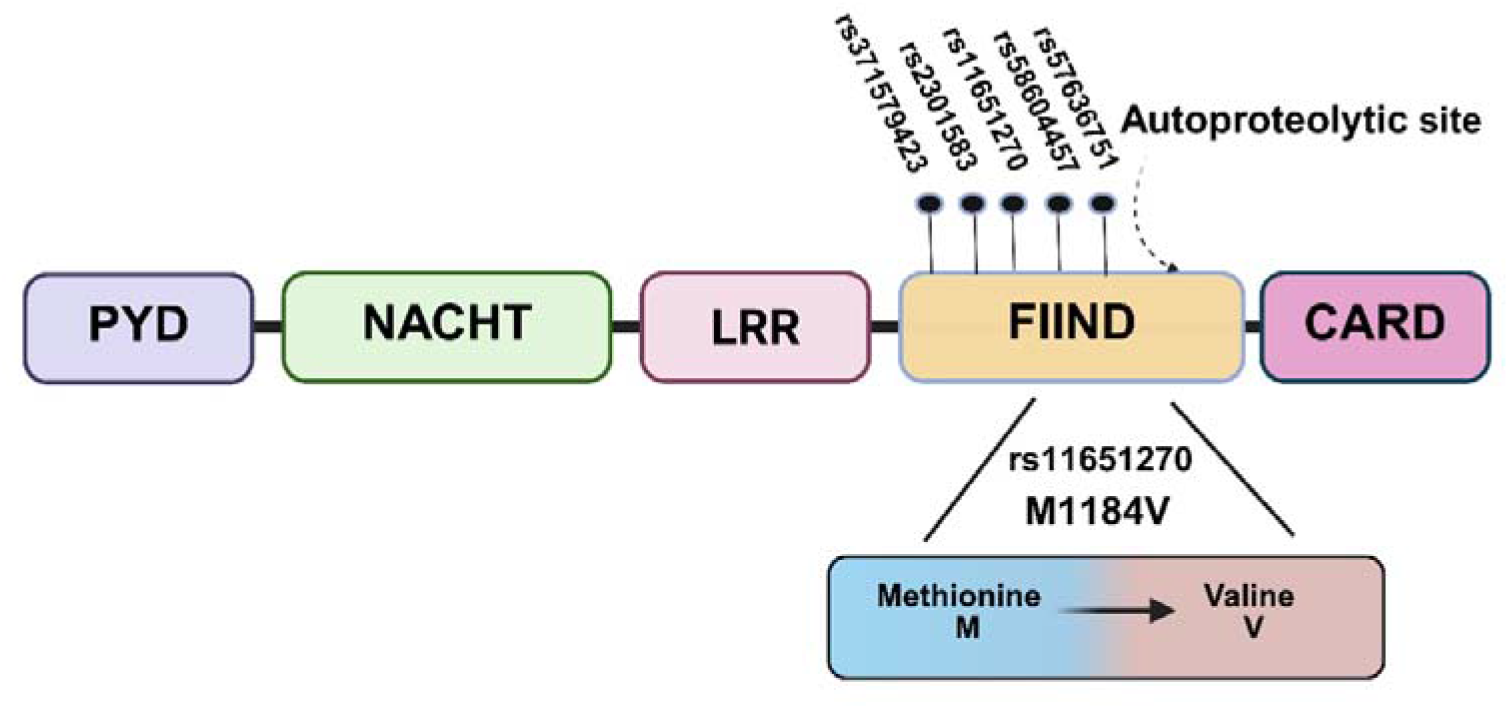
Schematic representation of *NLRP1* domain organization and FIIND-domain variants: *NLRP1* showing the pyrin domain (PYD), NACHT domain, leucine-rich repeat (LRR), function-to-find domain (FIIND), and caspase recruitment domain (CARD). Multiple variants identified in the FIIND region are indicated, including rs11651270, which results in a methionine-to-valine substitution at residue 1184.

### 2. Linkage Disequilibrium and Hardy–Weinberg Equilibrium Analysis of *NLRP1* Variants

Pairwise linkage disequilibrium analysis of all SNPs was performed on the Rajasthan control cohort (N = 756), and it revealed that rs58604457 and rs57636751 are in perfect LD (r^2^ = 1.0 and D’ = 1.0) with an identical MAF of 0.160, indicating that these variants represent the same genetic signal and cannot be treated as independent loci. The remaining variants displayed low-to-moderate LD: rs11651270 (r^2^ = 0.27, D′ = 0.97) and rs371579423 (r^2^ = 0.03, D′ = 0.715). Surprisingly, rs371579423 showed negligible allelic correlation with all other variants (r^2^ = 0.003–0.020), which confirms its statistical independence despite an elevated D’ (0.715) value. The four analysed SNPs corresponded to three LD-defined independent association signals: rs371579423, rs11651270, and the rs58604457/rs57636751 pair (Supplementary Table S2).

### 3. Genetic association analysis of *NLRP1* FIIND domain variants with glioma susceptibility

To evaluate whether polymorphisms in the FIIND domain of *NLRP1* contribute to glioma susceptibility, dominant, recessive, and codominant models were performed on four SNPs. Allelic and genotypic association statistics were calculated, and adjusted using a Bonferroni correction was applied across the three LD-defined independent association signals.

The rare variant rs371579423 (C>T; pos: 5521426) showed no evidence of association with glioma susceptibility. The MAF was nearly identical in cases (2.9%) and controls (3.0%). Allelic association testing across multiple case: control ratios (1:1, 1:2, and 1:10.8) revealed no significant association with glioma risk (N=756: p = 1.000; N=140: p = 0.782; N=70: p = 0.541). Estimated effect sizes were close to the null (OR = 0.72-0.96), with wide confidence intervals reflecting the low MAF of the variant. Similarly, the common coding-region SNP rs11651270 (T>C), which results in an M1184V substitution in the FIIND domain, showed no statistically significant association with glioma risk. In the primary analysis (70 cases vs. 756 controls), the alternate C allele frequency was 45.0% in cases and 40.0% in controls (OR = 1.23, 95% CI 0.85-1.76, p = 0.280), showing nearly matching results. Furthermore, 1:2 and 1:1 subsets (1:2 subset OR = 1.09, p = 0.677; 1:1 subset OR = 1.43, p = 0.181) demonstrated consistent, non-significant results and agreed with the primary analysis. Although ORs were > 1 in all three cases, even with the higher C allele frequency among cases, none reached nominal or corrected significance.

**Table 1:**
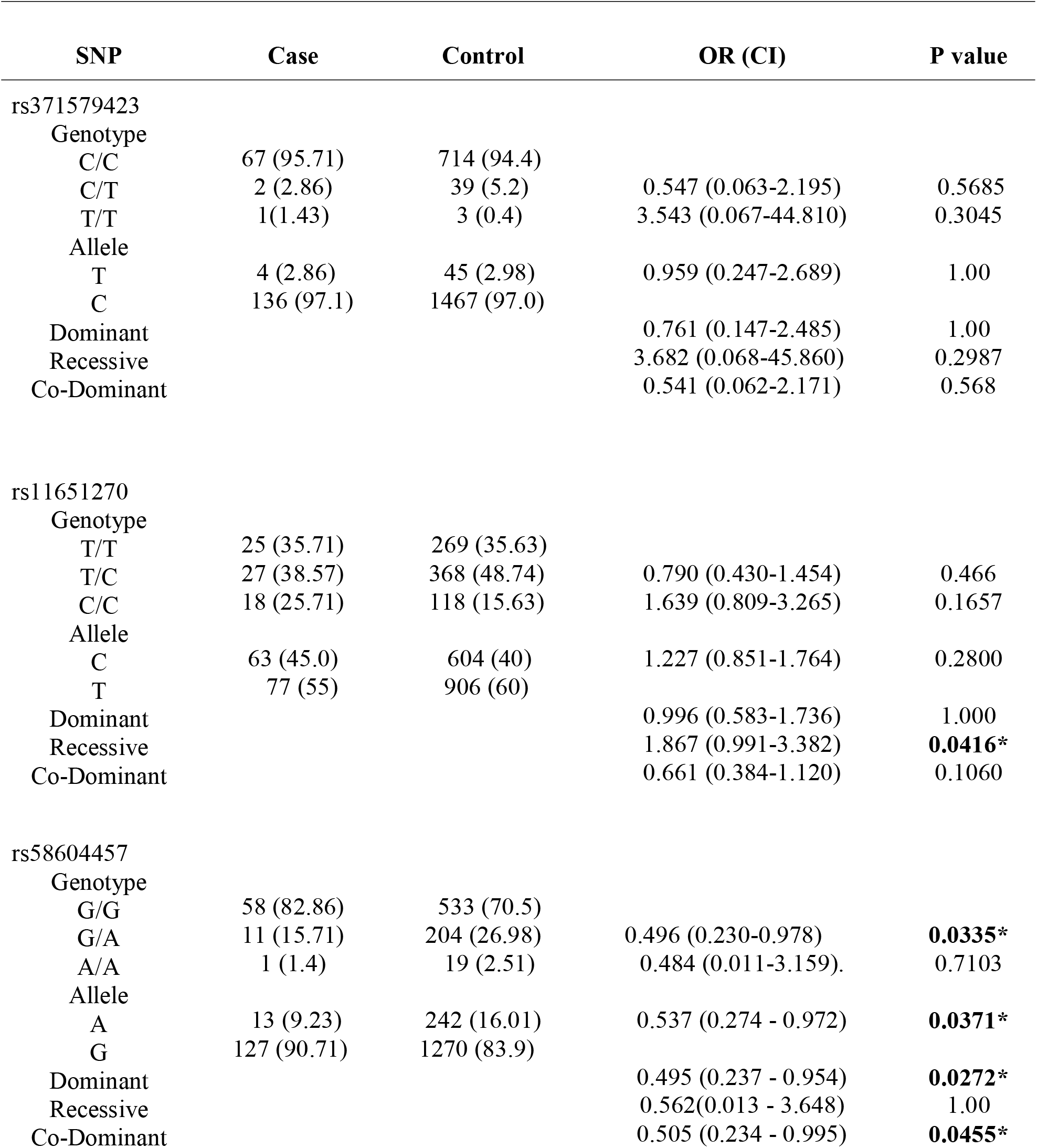
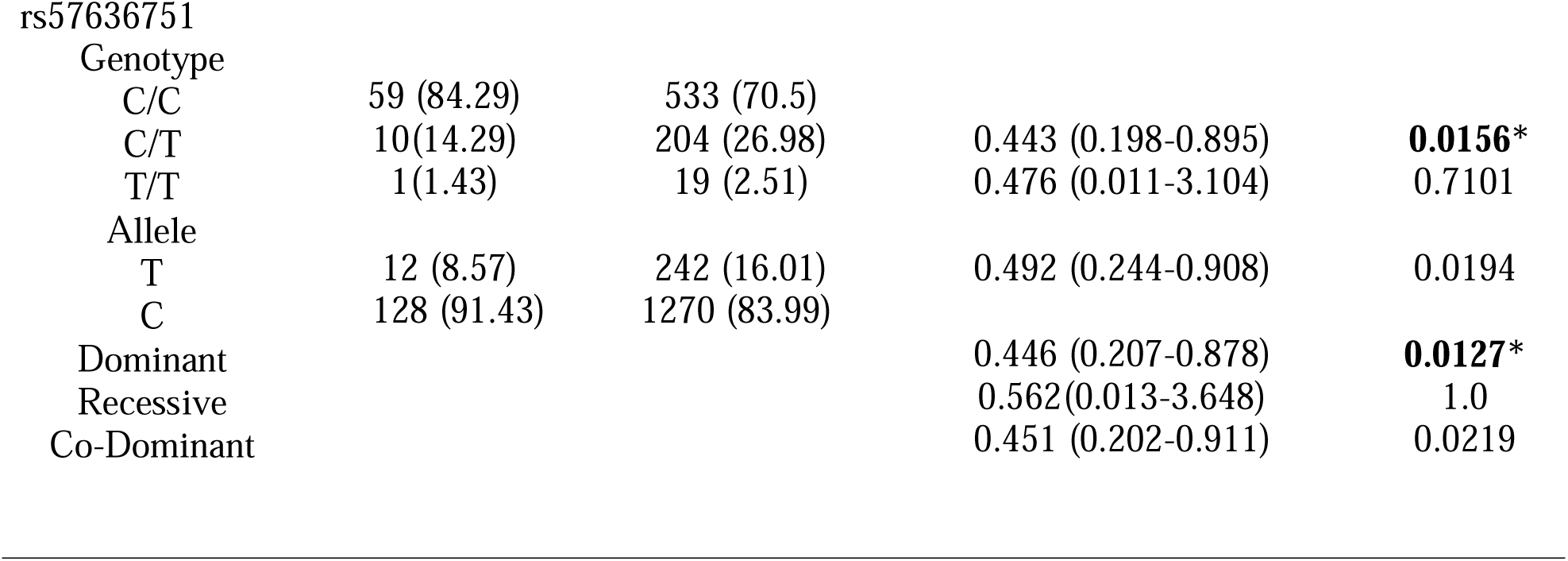
Genotype Frequency Distribution and association of variants across the Study Cohort.

| SNP | Case | Control | OR (CI) | P value |
| --- | --- | --- | --- | --- |
| rs371579423 |  |  |  |  |
| Genotype |  |  |  |  |
| C/C | 67 (95.71) | 714 (94.4) |  |  |
| C/T | 2 (2.86) | 39 (5.2) | 0.547 (0.063-2.195) | 0.5685 |
| T/T | 1(1.43) | 3 (0.4) | 3.543 (0.067-44.810) | 0.3045 |
| Allele |  |  |  |  |
| T | 4 (2.86) | 45 (2.98) | 0.959 (0.247-2.689) | 1.00 |
| C | 136 (97.1) | 1467 (97.0) |  |  |
| Dominant |  |  | 0.761 (0.147-2.485) | 1.00 |
| Recessive |  |  | 3.682 (0.068-45.860) | 0.2987 |
| Co-Dominant |  |  | 0.541 (0.062-2.171) | 0.568 |
| rs11651270 |  |  |  |  |
| Genotype |  |  |  |  |
| T/T | 25 (35.71) | 269 (35.63) |  |  |
| T/C | 27 (38.57) | 368 (48.74) | 0.790 (0.430-1.454) | 0.466 |
| C/C | 18 (25.71) | 118 (15.63) | 1.639 (0.809-3.265) | 0.1657 |
| Allele |  |  |  |  |
| C | 63 (45.0) | 604 (40) | 1.227 (0.851-1.764) | 0.2800 |
| T | 77 (55) | 906 (60) |  |  |
| Dominant |  |  | 0.996 (0.583-1.736) | 1.000 |
| Recessive |  |  | 1.867 (0.991-3.382) | **0.0416\*** |
| Co-Dominant |  |  | 0.661 (0.384-1.120) | 0.1060 |
| rs58604457 |  |  |  |  |
| Genotype |  |  |  |  |
| G/G | 58 (82.86) | 533 (70.5) |  |  |
| G/A | 11 (15.71) | 204 (26.98) | 0.496 (0.230-0.978) | **0.0335\*** |
| A/A | 1 (1.4) | 19 (2.51) | 0.484 (0.011-3.159). | 0.7103 |
| Allele |  |  |  |  |
| A | 13 (9.23) | 242 (16.01) | 0.537 (0.274 - 0.972) | **0.0371\*** |
| G | 127 (90.71) | 1270 (83.9) |  |  |
| Dominant |  |  | 0.495 (0.237 - 0.954) | **0.0272\*** |
| Recessive |  |  | 0.562(0.013 - 3.648) | 1.00 |
| Co-Dominant |  |  | 0.505 (0.234 - 0.995) | **0.0455\*** |
| rs57636751 |  |  |  |  |
| Genotype |  |  |  |  |
| C/C | 59 (84.29) | 533 (70.5) |  |  |
| C/T | 10(14.29) | 204 (26.98) | 0.443 (0.198-0.895) | **0.0156\*** |
| T/T | 1(1.43) | 19 (2.51) | 0.476 (0.011-3.104) | 0.7101 |
| Allele |  |  |  |  |
| T | 12 (8.57) | 242 (16.01) | 0.492 (0.244-0.908) | 0.0194 |
| C | 128 (91.43) | 1270 (83.99) |  |  |
| Dominant |  |  | 0.446 (0.207-0.878) | **0.0127\*** |
| Recessive |  |  | 0.562(0.013-3.648) | 1.0 |
| Co-Dominant |  |  | 0.451 (0.202-0.911) | 0.0219 |

The other two FIIND domain variants, viz., rs58604457 (G>A; pos: 5521873) and rs57636751 (C>T; pos: 5521934), demonstrated a protective association. LD analysis of controls showed that they represent a single underlying association signal (r^2^ = 1.00, D′ = 1.00). The A alternate allele of rs58604457 conferred a protective allelic effect (OR = 0.54, 95% CI 0.27–0.97, p = 0.037) as it occurred 1.7 times less frequently than in controls (9.3% vs 16.0%). Under a genotypic model, heterozygous G/A carriers exhibited a 50.0% reduction in glioma risk compared with G/G homozygotes (OR = 0.50, 95% CI 0.23–0.98, p = 0.034). Similarly, in rs57636751, the T allele frequency was underrepresented in cases, with an allelic OR of 0.49 (95% CI 0.24–0.91, p = 0.019). The heterozygous C/T individuals for the same SNP exhibited a 56.0% lower risk of developing glioma (OR = 0.44, 95% CI 0.20–0.90, p = 0.016) compared to C/C genotypic homozygosity. After Bonferroni correction across the three LD-defined independent association signals, the genotypic association for rs57636751 remained statistically significant (adjusted *p* = 0.047), whereas the other nominal associations did not exceed the corrected significance threshold.

**Fig. 2.**
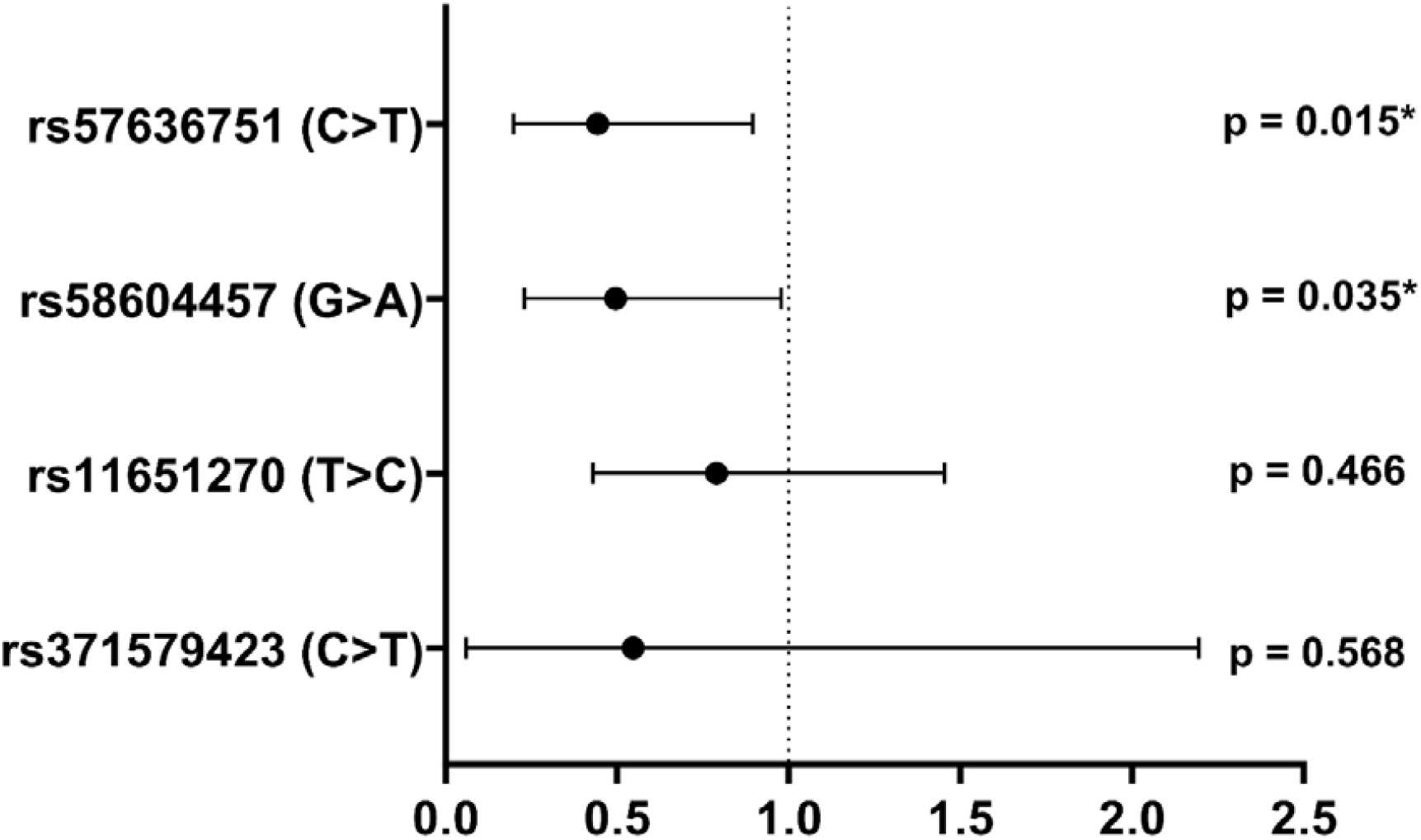
Forest plot of odds ratios for *NLRP1* FIIND domain variants in glioma. Odds ratios and 95% confidence intervals are shown for the minor allele of each SNP, comparing 70 Glioma cases against 756 Rajasthan population controls.

### 4. Distribution of *NLRP1* Variants Across Clinical and Demographic Parameters

To understand the variant distribution with respect to gender, age, and grade, we studied the sex distribution among histologically confirmed glioma cases. The cohort was male-predominant (70.7%), suggesting that higher-grade gliomas are more frequent in males than females. This analysis agrees with the well-established epidemiological finding that gliomas occur more frequently in males[30].

**Fig. 3.**
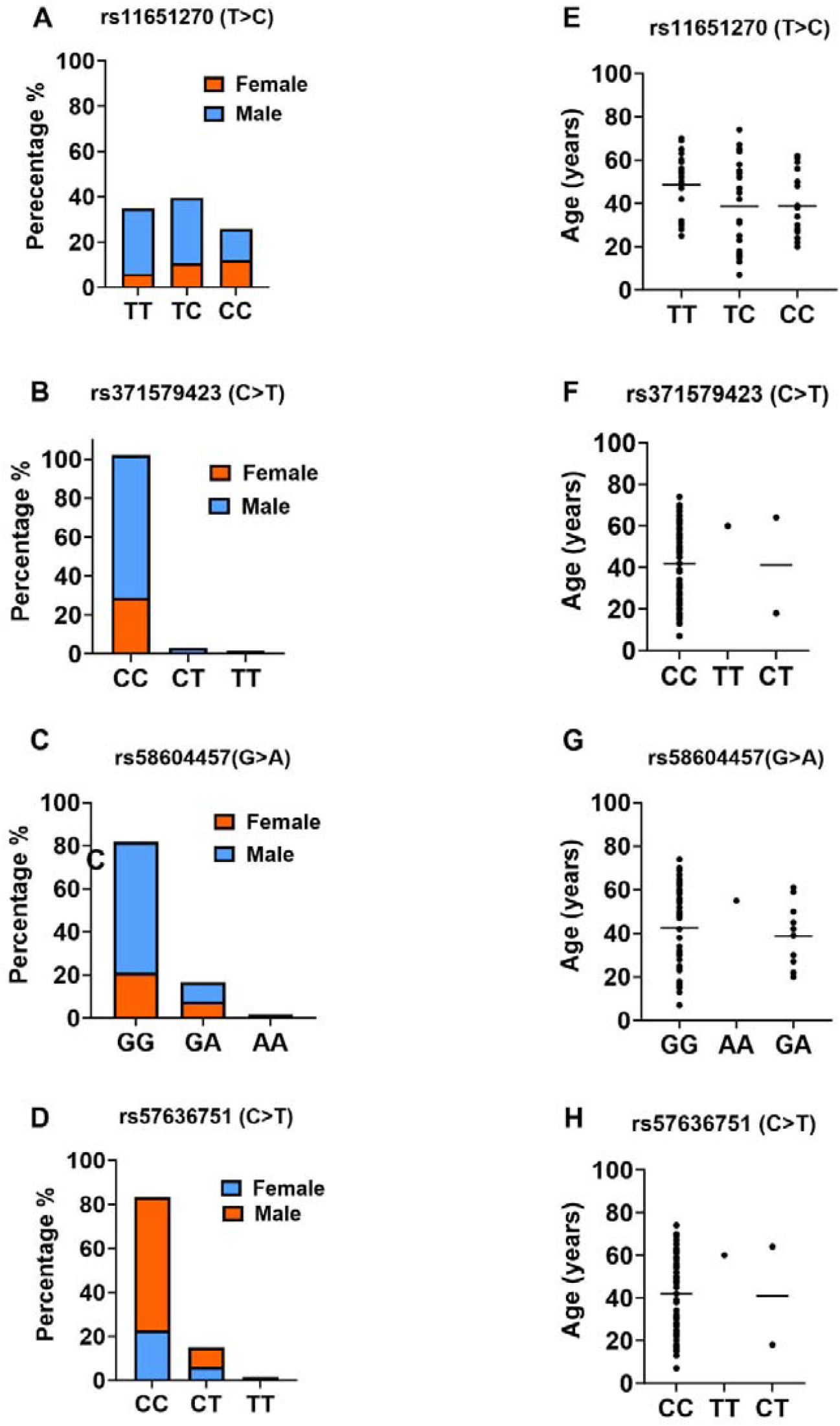
Gender distribution and age at diagnosis across *NLRP1* genotype groups in glioma cases (N=67). Left panels (A-D) show stacked bar charts depicting the percentage of male (blue) and female (orange) patients within each genotype group for four *NLRP1* variants. Right panels(E-H) show dot plots of age at diagnosis (years) stratified by genotype for each association between this variant and age of onset.

### 5. Structural and Molecular Dynamics Analysis of the *NLRP1* FIIND and CARD Domains and M1184V Variant

To examine the structural consequences of the M1184V substitution, we performed molecular dynamics simulations involving mutant and wild-type proteins in their pre- and post-cleavage states. MD simulations revealed interesting structural variability between wild-type and mutant systems as well as between pre- and post-cleavage states. As expected, the wild-type protein showed long-range excursions. It showed pronounced conformational rearrangements of the FIIND and CARD domains post-cleavage (Fig. 4), characterized by high backbone RMSD fluctuations (Fig 4b), an increased radius of gyration (Fig 4d), and large separation of the FIIND and CARD domains (Fig 4f). Interestingly, the mutant systems remained stable and structurally compact throughout the 100 ns simulation, maintaining lower RMSD values and minimal changes in the radius of gyration and inter-domain distances. Residue-level flexibility analysis revealed elevated fluctuations in specific regions of the wild-type cleaved structure (Fig S1), consistent with cleavage-induced structural plasticity, while the mutant displayed reduced mobility. Together, these results indicate that FIIND cleavage promotes large-scale conformational rearrangements in wild-type *NLRP1*, whereas the mutation was associated with reduced cleavage-associated conformational rearrangement in the simulated structure, which is stabilized.

**Fig. 4.**
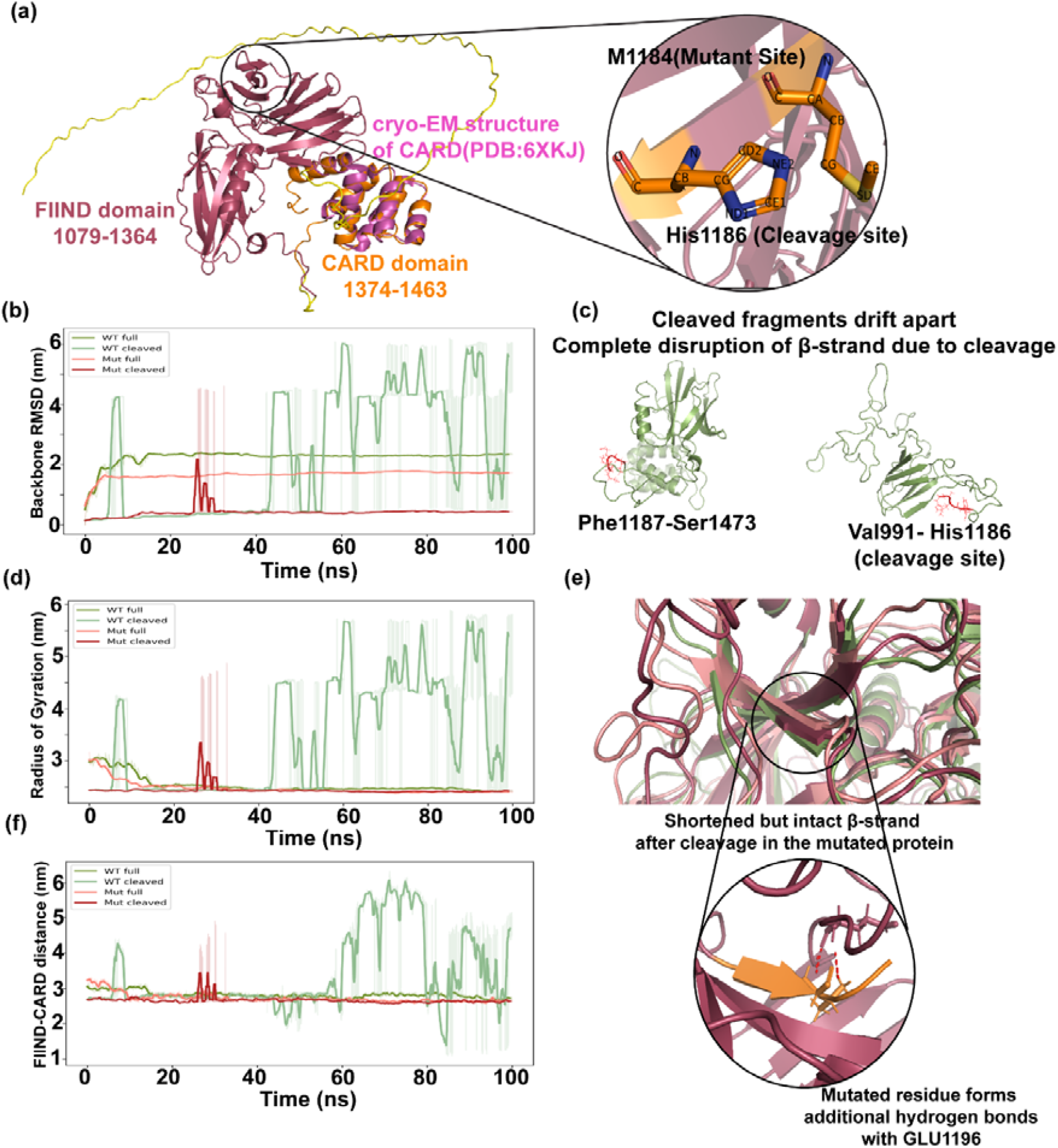
Structural and molecular representation of *NLRP1* FIIND and CARD domains. (A) AlphaFold2-predicted structure of FIIND and CARD domains, along with aligned cryo-EM structure of the CARD domain from PDB(6XKJ). Structural location of the mutant and cleavage sites within a β-strand(right). (B) Backbone RMSD showing flexibility of the structure throughout the simulation trajectory for wild-type and mutant proteins pre- and post-cleavage; (C) snapshot of the wild-type protein post-cleavage; (D) radius of gyration; (E) the mutated *nlrp1* retains a shorter beta strand even after cleavage, which is absent in the cleaved wild-type protein. The mutated residue forms hydrogen bonds with E1195. (F) Distance between the centre of mass of the FIIND and CARD domains.

The proximity of the mutant to the cleavage site is interesting to note, in addition to the fact that both sites are located within a - strand (Fig. 4a). On visual comparison of the final frames (taken from the simulation trajectory) of pre- and post-cleavage wild-type and mutated systems, we observed that the organization of residues into the - strand disappears in the post-cleavage state of the wild-type protein (Fig 4c). This disorientation appears to have been prevented by the M1184V mutant, where, despite cleavage, the length of the - strand seems to have shortened instead of being entirely disrupted and aligns well with the wild-type and mutant full-length protein (Fig 4e). This prompted us to look at the local nonpolar interactions of the mutated residue, where we find the formation of hydrogen bonds with E1195 from a neighbouring loop, which might prevent the structural disruption of the - strand and, consequently, the structural separation of the structural fragments on cleavage, in contrast to the wild-type protein. This hydrogen bond was found to be stably present throughout the simulation (>80% of frames), suggesting that this interaction may contribute to the local stability of the mutant structure after cleavage.

The total hydrogen bonds within residues gradually decrease in the cleaved wild-type protein as the simulation progresses on account of the cleaved fragments drifting away, as compared to the other systems, which maintain the domain architecture before and after cleavage, as indicated by the roughly similar number of hydrogen bonds throughout the trajectory. Furthermore, the mutant systems show comparatively less fluctuation in the number of hydrogen bonds than the full-length wild-type protein, indicating diminished local flexibility (Fig S1b).

## Discussion

In this pilot case-control study, we identified five *NLRP1* FIIND-domain variants in an Indian glioma cohort: rs371579423, rs58604457, rs57636751, rs11651270, and rs2301583. Two intronic variants, rs58604457 (G>A) and rs57636751(C>T), were in complete linkage disequilibrium (r^2^=1.00) and were associated with lower odds of glioma, whereas the coding variant rs11651270 (M1184V) showed no significant association with disease status. Under a dominant genotypic framework, the carriers of the minor alleles at this locus of interest exhibited a nominally significant protective association with glioma patients. The heterozygous C/T state of rs57636751 conferred a reduction in observed risk (OR = 0.443, 95% CI 0.198–0.895, p=0.0156) when compared to major-allele homozygotes. Following the application of a stringent Bonferroni correction for multiple hypothesis testing, this specific genotypic association maintained borderline statistical significance. ( 0.0468). As these variants are intronic, rather than coding, their potential functional consequences could involve transcript regulation or splicing; however, this possibility was not examined in the present study. These findings suggest that while common functional variants such as rs11651270 may not strongly influence glioma susceptibility in this cohort, other FIIND domain polymorphisms may contribute modest protective effects that suggest further investigation in larger populations. The rs2301583, which was not found in the GenomeIndia population-based catalogue, was annotated as benign in ClinVar and had a MAF of 6.0%. Its higher allelic frequency in the glioma cases makes it a candidate for further functional studies. The case cohort (N=70) limits statistical power to detect modest and/or rare-variant contributions and the observed odds ratios. Moreover, the borderline significance after multiple-testing correction for some variants underscores the need for validation in larger glioma-control cohorts.

To further investigate the potential structural consequences of FIIND domain variation, MD simulations were performed to compare the conformational behaviour of wild-type and M1184V mutant structures in both pre- and post-cleavage states, with particular emphasis on FIIND-CARD domain dynamics. Simulations indicate that the mutant protein remains structurally stable even after cleavage, consistent with earlier reports suggesting increased cleavage efficiency associated with this variant. However, the mechanism by which this mutation stabilizes the *NLRP1* structure remained unclear. We observed in our simulations that the mutant protein remains structurally stable following cleavage, whereas the wild-type protein loses β-strand organization at the cleavage site. This disruption appears to be limited in the mutant, where the β-strand shortens rather than fully unravels, coinciding with a stabilizing hydrogen bond formed between the mutated residue and E1195 on a neighbouring loop. Despite this local stabilization, the mutation did not substantially alter global FIIND-CARD domain dynamics across the simulation trajectory, suggesting the effect is confined to the immediate cleavage region rather than producing large-scale conformational rearrangement. The M1184V substitution was associated with preservation of local β-strand organization and formation of a persistent interaction with E1195 after cleavage, while global FIIND–CARD dynamics remained broadly comparable between mutant and wild-type trajectories. These findings suggest that the structural effect of M1184V is predominantly local rather than a large-scale rearrangement of the FIIND–CARD architecture. Together, these observations offer a plausible structural mechanism by which M1184V may stabilize the FIIND domain post-cleavage, with potential implications for *NLRP1* inflammasome regulation.

## Conclusions

This study provides a combined genetic and structural investigation of FIIND-domain variants of *NLRP1* in glioma in an Indian cohort. Statistical models showed that at least two variants are significantly associated with glioma susceptibility. However, rs11651270 did not show an association with glioma. Molecular dynamics simulations indicate that the M1184V substitution may stabilize local FIIND structural architecture following cleavage, potentially influencing *NLRP1* conformational dynamics independent of its lack of association with glioma risk. The modest sample size and use of population-based rather than individually matched controls are key limitations of this pilot study. These findings highlight the importance of integrating population genetics with structural and functional analyses to better understand the role of inflammasome-related genes in cancer biology. Future studies involving larger cohorts and functional validation will be necessary to clarify the contribution of *NLRP1* variation to glioma pathophysiology.

## Acknowledgement

The authors would like to thank Dr. Deepak Jha, AIIMS Jodhpur, India, for their support in providing glioma tissue samples. We thank the Technology Innovation and Start-up Centre (TISC), Jodhpur, for providing the Sanger sequencing facility. S.C. is thankful for the DST INSPIRE Fellowship; M.S.R., D.K., S.A., and J.S. are thankful for the MHRD fellowship.

## Author Contributions

**Shalini Chhipa:** Writing-original draft, Formal analysis, Data curation, investigation, and Methodology. **Omkar A. Sonawane:** Writing-original draft, Formal analysis, Investigation, and Methodology. **MS Revanth:** Writing-original draft, Formal analysis, Investigation, and Methodology. **Deepak Kumar:** Formal analysis, Data curation, Investigation. **Jyoti Sharma:** Formal analysis, investigation. **Srijita Acharya and Sucharita Dey:** Methodology and investigation. **Deepka Jha and Pankaj Yadav:** Resources. **Sushmita Jha:** Writing-conceptualization, review & editing, Project administration.

## Data Availability Statement

The data generated in this study are available within the article and its Supporting Information files.

## Ethics Approval and Consent to Participate

The Internal Review Board and the Ethics Committees of AIIMS, Jodhpur, approved the acquisition of glioma tissues. Human subjects gave their informed consent before tissue samples were used for research. Every experiment was carried out in compliance with the ethical standards and policies of the Indian Institute of Technology Jodhpur and AIIMS, Jodhpur.

