## supplementary table 1, 2, 3 for "Genetic and structural interpretation of NLRP1 FIIND domain variants in glioma in an Indian cohort: a pilot study"

**Supplementary Table 1: Hardy-Weinberg Equilibrium Test**

| **SNP** | **REF** | **ALT** | **Obs**  **REF_hom** | **Obs**  **het** | **Obs**  **ALT**  **hom** | **Exp.**  **Hom** | **Exp**  **het** | **Exp**  **ALT**  **hom** | **REF**  **AF** | **ALT**  **AF** | **Chi^2^** | **df** | **p**  **HWE** |
| --- | --- | --- | --- | --- | --- | --- | --- | --- | --- | --- | --- | --- | --- |
| rs371579423 | C | T | 714 | 39 | 3 | 711.67 | 43.66 | 0.67 | 0.9702 | 0.0298 | 8.6148 | 1 | 0.33 |
| rs11651270 | T | C | 269 | 368 | 118 | 271.8 | 362.4 | 120.8 | 0.6 | 0.4 | 0.1803 | 1 | 0.6711 |
| rs58604457 | G | A | 533 | 204 | 19 | 533.37 | 203.27 | 19.37 | 0.8399 | 0.1601 | 0.0098 | 1 | 0.921 |
| rs57636751 | C | T | 533 | 204 | 19 | 533.37 | 203.27 | 19.37 | 0.8399 | 0.1601 | 0.0098 | 1 | 0.921 |

**Supplementary Table 2: Linkage Disequilibrium in Control samples**

| **SNP1** | **POS1** | **SNP2** | **POS2** | **r^2^** | **D_prime** | **AF1** | **AF2** | **LD_class** |
| --- | --- | --- | --- | --- | --- | --- | --- | --- |
| rs371579423 | 5521426 | rs11651270 | 5521757 | 0.0196 | 0.989 | 0.0291 | 0.4 | low |
| rs371579423 | 5521426 | rs58604457 | 5521873 | 0.003 | 0.7146 | 0.0298 | 0.1601 | low |
| rs371579423 | 5521426 | rs57636751 | 5521934 | 0.003 | 0.7146 | 0.0298 | 0.1601 | low |
| rs11651270 | 5521757 | rs58604457 | 5521873 | 0.2729 | 0.9764 | 0.4 | 0.1603 | low |
| rs11651270 | 5521757 | rs57636751 | 5521934 | 0.2729 | 0.9764 | 0.4 | 0.1603 | low |
| rs58604457 | 5521873 | rs57636751 | 5521934 | 1 | 1 | 0.1601 | 0.1601 | Pefect LD |

**Supplementary Table 3: rsIDs and respective MAF**

| **rsID** | **Ref** | **Alt** | **VEP**  **Consequence** | **VEP**  **Impact** | **Global_MAF** | **Cohort_MAF** |
| --- | --- | --- | --- | --- | --- | --- |
| rs371579423 | C | T | intron_variant | MODIFIER | 0.001592 | 0.0063 |
| rs2301583&COSV52581359 | G | A | synonymous_variant | LOW | 0.01372 | 0.0625 |
| rs11651270&CM1212937&COSV52564879 | T | C | missense_variant | MODERATE | 0.4553 | 0.2245 |
| rs58604457 | G | A | intron_variant | MODIFIER | 0.03259 | 0.0548 |
| rs57636751 | C | T | intron_variant | MODIFIER | 0.03935 | 0.0541 |

**Supplementary Figure 1 (S1)**

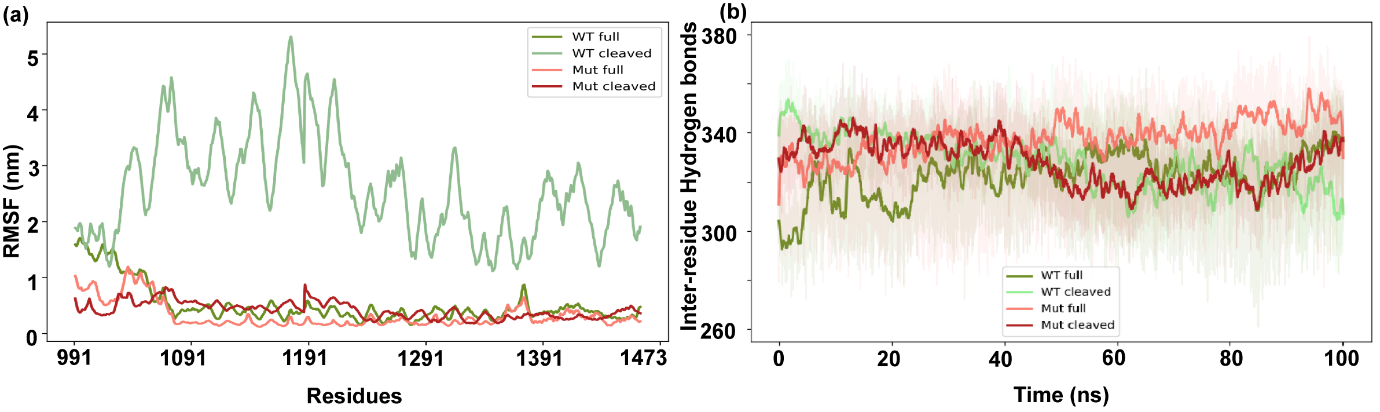

Fig S1. (a) Residue-wise flexibility of the NLRP1 FIIND and CARD domains.(b) Fluctuation of hydrogen bonding between residues throughout the simulation time.
